# Genomic epidemiology revealed the emergence and worldwide dissemination of ST383 carbapenem-resistant hypervirulent *Klebsiella pneumoniae* and hospital acquired infections of ST196 *Klebsiella quasipneumoniae* in Qatar

**DOI:** 10.1101/2022.06.02.494628

**Authors:** Clement Kin-Ming Tsui, Fatma Ben Abid, Christi Lee McElheny, Muna Almuslamani, Ali S. Omrani, Yohei Doi

**Author notes:** Address correspondence to Clement Tsui, and Fatma B. Abid. National Centre for Infectious Diseases, Tan Tock Seng Hospital, Singapore (Clement Kin-Ming Tsui).

## Abstract

The emergence of carbapenem-resistant (CR) hypervirulent *Klebsiella pneumoniae* (hvKp) is a new threat to healthcare. In this study, we studied the molecular epidemiology of CR *Klebsiella* isolates in Qatar using whole genome sequence data. We also characterised the prevalence and genetic basis of hypervirulent phenotypes, and established the virulence potential using a *Galleria mellonella* model. One hundred CR *Klebsiella* isolates were recovered, and NDM and OXA-48 were the most common carbapenemases. Phylogenetic analysis indicated the presence of diverse sequence types and clonal lineages; one of them belonged to *K. quaisipneumoniae* ST196 that may be disseminated among several health care centres. Ten *K. pneumoniae* isolates carrying *rmpA* and/or *rmpA2*, and 2 isolates belonged to KL2, indicating the prevalence of classical hypervirulent (hv) isolates was not high. Isolates carrying CR and hv genes were confined mainly to ST231 and ST383 isolates. One ST383 isolate was further investigated by MinION sequencing, and the assembled genome indicated the *bla*_NDM_ was located on an IncHI1B type plasmid (pFQ61_ST383_NDM-5), which also harbored several virulence factors, including the regulator of the mucoid phenotype (*rmpA*), the regulator of mucoid phenotype 2 (*rmpA2*), and aerobactin (*iucABCD* and *iutA*), likely resulting from inversion and recombination events. In contrast, *bla*_OXA-48_ was located in an IncL-type plasmid. Comparative genomes indicated the recent evolution and emergence of CR-hv Kp ST383 via the acquisition of hybrid plasmids with both carbapenemase and virulence genes. CR-hv *K. pneumoniae* ST383 pose an emerging threat to global health due to their simultaneous hypervirulence and multidrug resistance.

## Introduction

*Klebsiella pneumoniae* are Gram-negative bacterial pathogens that are widely present in nature and in the human intestine. *K. pneumoniae* are well known to cause hospital-acquired infections in immunocompromised patients (1, 2), but infections caused by *K. pneumoniae* can also occur in long-term care facilities, such as nursing homes, and in the community. Types of infection vary and include hospital-acquired pneumonia, lung abscesses, bacteremia, catheter-related infections, wound or surgical site infections, upper and lower urinary tract infections, liver abscesses, and meningitis (1). Based on the molecular and genome sequencing data, various novel species and subspecies, such as *K. michiganensis, K. quasipneumoniae*, and *K. variicola*, have been recognized(3). In our previous study, where we had studied 149 CRE isolates in Qatar, *K. pneumoniae* and *K. quasipneumoniae* isolates were prevalent (4); over 50% of the collections belonged to *K. pneumoniae*.

Carbapenem-resistant *Enterobacterales* (CRE) infections are a global health priority and are among the most serious antimicrobial resistance (AMR) threats (5). *K. pneumoniae* is the most common species among CRE. Carbapenem resistance in *K. pneumoniae* is primarily driven by production of carbapenemases, with extended spectrum beta-lactamases (ESBL) playing a supplementary role (4, 6–9). In the aforementioned study, genes encoding metallo-β-lactamases were detected in 45.8% of the isolates, and OXA-48-like enzymes in 40.3%.

Hypervirulent *K. pneumoniae* (hvKp) can cause life-threatening, community-acquired infections such as liver abscesses, and are associated with high mortality and morbidity (10). Several virulence factors contributed to the pathogenicity, including hypermucoviscous-specific capsular antigens (i.e., K1 and K2 serotypes) and virulence genes such as *rmpA* (regulator of mucoid phenotype) and aerobactin (10, 11). Traditionally, multidrug-resistant (MDR) and hypervirulent (hv) phenotypes in *K. pneumoniae* were associated with distinct lineages. However, MDR lineages acquiring virulence traits or hypervirulent lineages acquiring resistance genes have increasingly been reported in the last decade, especially in South and South-East Asia (12–15). Also, hvKp have been evolving and becoming more resistant than the classical *K. pneumoniae* strain(16), mostly through the dissemination of conjugative and hybrid plasmids. This may lead to widely disseminated community-acquired infections in healthy people that are difficult to treat.

While carbapenem resistance in *K. pneumoniae* is increasingly documented in the Middle East region, there is limited information on hypervirulence and how it intersects with carbapenem resistance. We therefore conducted an in-depth analysis of *Klebsiella* genomes that were sequenced in the study of CRE in Qatar, with the following three aims: (i) to describe the genetic diversity, AMR and virulence determinants of *Klebsiella* isolates; (ii) to relate the molecular epidemiology data of selected sequence types (ST) to observations made in other parts of the world, and (iii) to characterize the genetic context and virulence potential of a carbapenemase-resistant and hypervirulent *Klebsiella pneumoniae* (CR-hvKp) isolate.

## Results

### Epidemiology of sequence types and AMR genes

*Klebsiella pneumoniae* (Kp) was the most common species (n=80), followed by *K. quasipneumoniae* (n=16), *K. aerogenes* (n=3) and *K. oxytoca* (n=1). Over 40 different STs were reported, and 23 (28.8%) *K. pneumoniae* STs were represented by a single isolate. *K. pneumoniae* ST147 was predominant (n=13), followed by ST231 (n=7) and ST11 (n=5). ST147 and ST11 belong to widespread clonal groups (CG) 147 and CG258, while the ‘traditional’ high risk CG14/15 (n=4) was not common, and CC101 and CC17 were not found (Table 1, Table S1). Nine *K. quasipneumoniae* isolates belonged to ST196 and four belonged to ST1416, and these STs belonged to *K. quasipneumoniae* subsp. *quasipneumoniae*. Across different specimen types, Kp ST147 was isolated from all specimens, followed by ST231. Similarly, *K. quasipneumoniae* subsp. *quasipneumoniae* can be isolated from blood, pus and urine (Table 1).

**Table 1.**
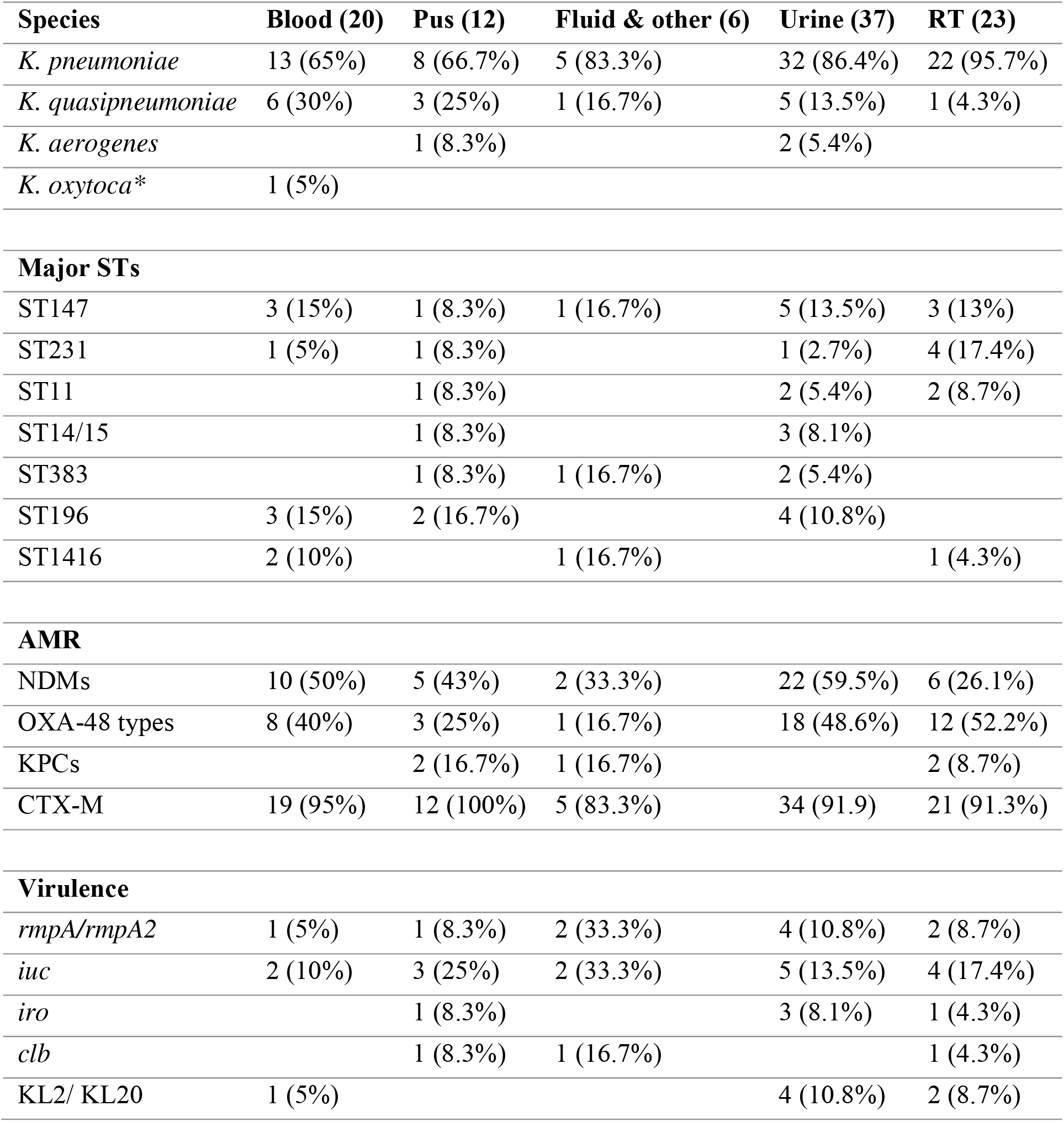
Comparison of key features of *Klebsiella* samples from different specimens *(as *K. michiganensis* in Kleborate).

| Species | Blood (20) | Pus (12) | Fluid & other (6) | Urine (37) | RT (23) |
| --- | --- | --- | --- | --- | --- |
| *K. pneumoniae* | 13 (65%) | 8 (66.7%) | 5 (83.3%) | 32 (86.4%) | 22 (95.7%) |
| *K. quasipneumoniae* | 6 (30%) | 3 (25%) | 1 (16.7%) | 5 (13.5%) | 1 (4.3%) |
| *K. aerogenes* |  | 1 (8.3%) |  | 2 (5.4%) |  |
| *K. oxytoca* * | 1 (5%) |  |  |  |  |
| **Major STs** |  |  |  |  |  |
| ST147 | 3 (15%) | 1 (8.3%) | 1 (16.7%) | 5 (13.5%) | 3 (13%) |
| ST231 | 1 (5%) | 1 (8.3%) |  | 1 (2.7%) | 4 (17.4%) |
| ST11 |  | 1 (8.3%) |  | 2 (5.4%) | 2 (8.7%) |
| ST14/15 |  | 1 (8.3%) |  | 3 (8.1%) |  |
| ST383 |  | 1 (8.3%) | 1 (16.7%) | 2 (5.4%) |  |
| ST196 | 3 (15%) | 2 (16.7%) |  | 4 (10.8%) |  |
| ST1416 | 2 (10%) |  | 1 (16.7%) |  | 1 (4.3%) |
| **AMR** |  |  |  |  |  |
| NDMs | 10 (50%) | 5 (43%) | 2 (33.3%) | 22 (59.5%) | 6 (26.1%) |
| OXA-48 types | 8 (40%) | 3 (25%) | 1 (16.7%) | 18 (48.6%) | 12 (52.2%) |
| KPCs |  | 2 (16.7%) | 1 (16.7%) |  | 2 (8.7%) |
| CTX-M | 19 (95%) | 12 (100%) | 5 (83.3%) | 34 (91.9) | 21 (91.3%) |
| **Virulence** |  |  |  |  |  |
| *rmpA/rmpA2* | 1 (5%) | 1 (8.3%) | 2 (33.3%) | 4 (10.8%) | 2 (8.7%) |
| *iuc* | 2 (10%) | 3 (25%) | 2 (33.3%) | 5 (13.5%) | 4 (17.4%) |
| *iro* |  | 1 (8.3%) |  | 3 (8.1%) | 1 (4.3%) |
| *clb* |  | 1 (8.3%) | 1 (16.7%) |  | 1 (4.3%) |
| KL2/ KL20 | 1 (5%) |  |  | 4 (10.8%) | 2 (8.7%) |

The most prevalent carbapenemase genes were those encoding NDM-1 (n=39), OXA-48 (n=20), OXA-232 (n=10) and OXA-181 (n=12). KPC-2 (n=3) and KPC-3 (n=2) were identified but rare. Fifteen isolates did not carry carbapenemase genes, but carried combinations of *bla*_CTX-M_ genes and loss-of-function genetic changes in *ompK35* and *ompK36* (Table S1), which has previously been linked to carbapenem resistance in *Klebsiella* (17). In total, 68 out of 90 *Klebsiella* samples had *ompK35* or *ompK36* loss/truncations/mutations, which may contribute to reduced susceptibility to carbapenems (Table S1). Genes encoding CTX-M-type ESBL were co-harbored by 75 isolates (75/85, 88.2%), including 66 isolates (77.6%) harboring *bla*_CTX-M-15_ genes, 7 isolates (8.2%) harboring *bla*_CTX-M-14b_ genes, and 2 isolates (2.3%) harboring *bla*_CTX-M-27_ genes. Co-carriage of NDM- and/or OXA-48 type carbapenemase genes and a CTX-M gene was reported in 4 isolates, including three *K. pneumoniae* ST383 isolates. *bla*_OKP_, an intrinsic beta-latamase gene in *K. quasipneumoniae*, 13 was found to be uniquely restricted within the *K. quasipneumoniae* cluster. Specifically, blaOKP-A was associated with *K. quasipneumoniae* subsp. *quasipneumoniae* and blaOKP-B with *K. quasipneumoniae* subsp. *similipneumoniae* (Table S1).

Phylogenetic analysis based on core genome SNPs illustrated that *K. pneumoniae* and *K. quasipneumoniae* isolates formed monophyletic clades respectively. There were variations within the prevalent *K. pneumoniae* STs with respect to the genetic distance and presence/absence of certain AMR genes such as ST307 and ST147; in contrast, *K. quasipneumoniae* var. *quasipneumoniae* ST196 was highly clonal (Figure 1).

**Figure 1.**
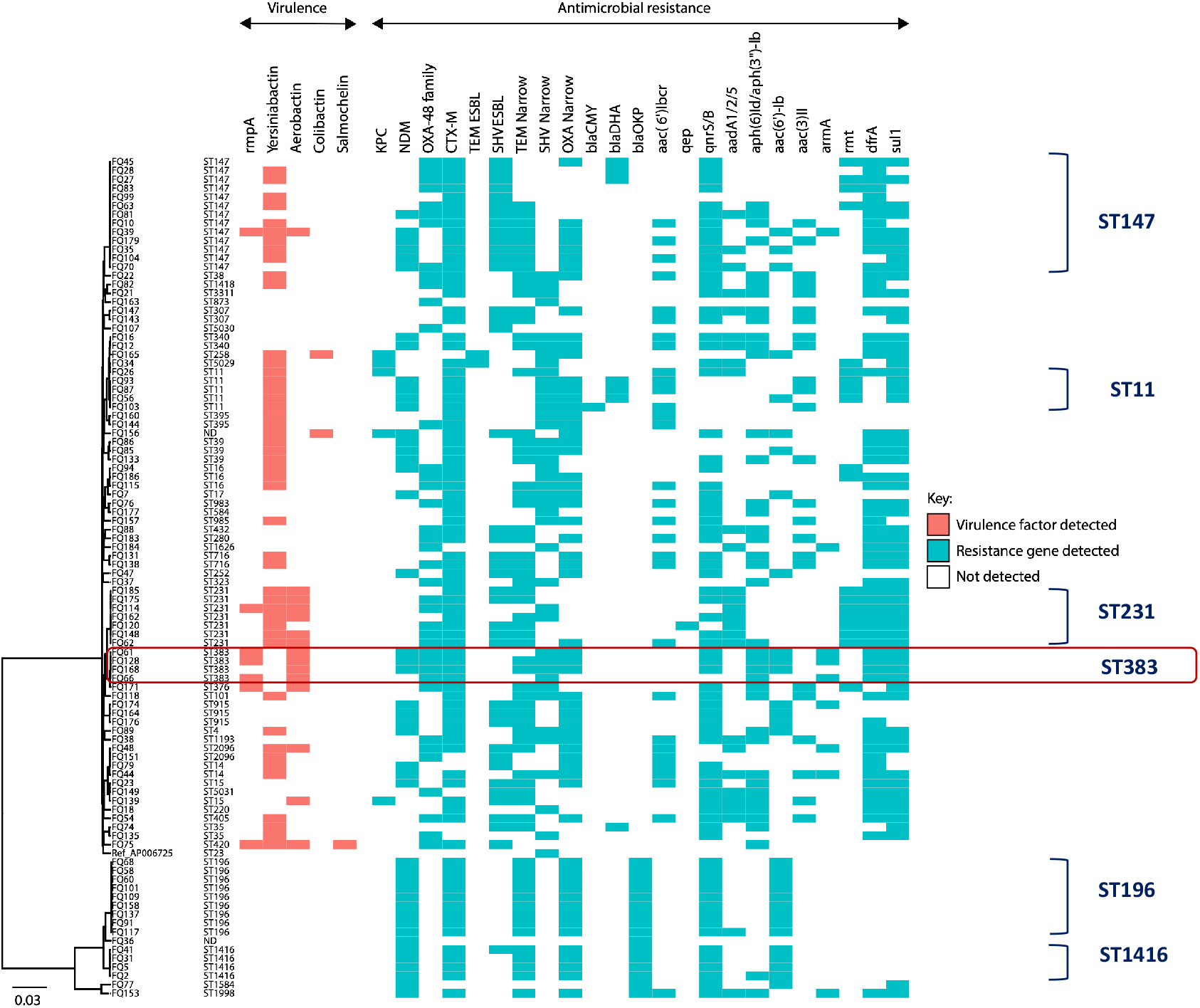
Genetic relationship of *K. pneumoniae* and *K. quasipneumoniae* isolates inferred from core genome SNPs overlaid with the presence/absence of antimicrobial resistance and virulence phenotypes.

Plasmid location was inferred for 24 carbapenemase genes by Plasmidfinder on short read assemblies; *bla*_NDM-1_ was localized on IncFII replicon, while *bla*_OXA-181_ and *bla*_OXA-232_ were commonly associated with ColKP3 plasmids (Table 2). A pOXA-48-like plasmid was detected in 17/20 isolates harbouring *bla*_OXA-48_. In contrast, most contigs carrying *bla*_NDM_ were divergent and had different mobile genetic elements. While NDM-1 was linked to IncFIB and IncA/C2 in *K. pneumoniae, bla*_NDM-1_ in seven out of nine *K. quasipneumoniae* ST196 isolates were found to be associated with IncFII_1_pKP91. Indeed, the contig carrying *bla*_NDM-1_ and *bla*_CTX-M-15_ was homologous to that in *K. quasipneumoniae* ST1416, suggesting a plasmid may be circulating and conferring carbapenem resistance to other STs. *bla*_KPC-3_ was located on a contig with the commonly reported mobile genetic element Tn*4401a* (Figure S1).

**Table 2.** Plasmid location of carbapenemase genes in 24 isolates - *K. pneumoniae* and *K. quasipneumoniae* (from Plasmidfinder)

| Sample ID # | Species | ST | Date | Carbapenemase | Plasmid |
| --- | --- | --- | --- | --- | --- |
| FQ103 | *K. pneumoniae* | 11 | 16.03.17 | **NDM-1** | **IncA/C2** |
| FQ7 | *K. pneumoniae* | 17 | 20.05.15 | **NDM-1** | **IncFIB(pQil)** |
| FQ94 | *K. pneumoniae* | 16 | 26.02.17 | **OXA-181** | **ColKP3** |
| FQ115 | *K. pneumoniae* | 16 | 27.04.17 | **OXA-181** | **ColKP3** |
| FQ186 | *K. pneumoniae* | 16 | 24.12.17 | **OXA-232** | **ColKP3** |
| FQ22 | *K. pneumoniae* | 38 | 07.07.15 | **OXA-232** | **ColKP3** |
| FQ27 | *K pneumoniae* | 147 | 22.07.15 | **OXA-181** | **ColKP3** |
| FQ28 | *K. pneumoniae* | 147 | 25.07.15 | **OXA-181** | **ColKP3** |
| FQ45 | *K. pneumoniae* | 147 | 11.11.15 | **OXA-181** | **ColKP3** |
| FQ70 | *K. pneumoniae* | 147 | 21.10.16 | **OXA-181** | **ColKP3** |
| FQ114 | *K. pneumoniae* | 231 | 25.04.17 | **OXA-232** | **ColKP3** |
| FQ120 | *K. pneumoniae* | 231 | 17.05.17 | **OXA-232** | **ColKP3** |
| FQ148 | *K. pneumoniae* | 231 | 26.08.17 | **OXA-232** | **ColKP3** |
| FQ144 | *K. pneumoniae* | 395 | 20.08.17 | **OXA-232** | **ColKP3** |
| FQ138 | *K. pneumoniae* | 716 | 27.07.17 | **OXA-48** | **ColKP3** |
| FQ151 | *K. pneumoniae* | 2096 | 05.09.17 | **OXA-232** | **ColKP3** |
| FQ107 | *K. pneumoniae* | 5030 | 26.03.17 | **OXA-232** | **ColKP3** |
| FQ149 | *K pneumoniae* | 5031 | 28.08.17 | **OXA-181** | **ColKP3** |
| FQ58 | *K. quasipneumoniae subsp. quasipneumoniae* | 196 | 03.04.16 | **NDM-1** | **IncFII_1_pKP91** |
| FQ60 | *K. quasipneumoniae subsp. quasipneumoniae* | 196 | 22.04.16 | **NDM-1** | **IncFII_1_pKP91** |
| FQ91 | *K. quasipneumoniae subsp. quasipneumoniae* | 196 | 12.02.17 | **NDM-1** | **IncFII_1_pKP91** |
| FQ101 | *K. quasipneumoniae subsp. quasipneumoniae* | 196 | 14.03.17 | **NDM-1** | **IncFII_1_pKP91** |
| FQ117 | *K. quasipneumoniae subsp. quasipneumoniae* | 196 | 28.04.17 | **NDM-1** | **IncFII_1_pKP91** |
| FQ137 | *K. quasipneumoniae subsp. quasipneumoniae* | 196 | 26.07.17 | **NDM-1** | **IncFII_1_pKP91** |

**Table 2a.** Notable hypervirulent isolates in this investigation (virulence markers from Kleborate). RT: respiratory tract; UTI: Urinary tract infection; IAI: Intrabdominal infection; BSI: Blood stream infection; RTI: Respiratory infection; SSTI: Skin & soft tissue infection; Yersiniabactin (ybt); Aerobactin (iuc); Salmochelin (iro); Colibactin (clb); ESBL: extended spectrum beta-lactamase

| ID # | Specimen | Infection | ST | *rmpA* | *rmpA2* | *ybt* | *iuc* | *iro* | *clb* | *K_locus* | Carbapenemases | ESBL | Other ESBL | Omp mutations | Clinical outcome | Travel (30 days) |
| --- | --- | --- | --- | --- | --- | --- | --- | --- | --- | --- | --- | --- | --- | --- | --- | --- |
| FQ44 | Urine | UTI | 14 | - | - | ybt 14; ICEKp5 |  |  |  | KL2 | NDM-1 | CTX-M-15 | - | OmpK35-88%; OmpK36GD | alive | no |
| FQ139 | Pus | IAI | 15 | - | - | - | iuc 5 |  |  | KL102 | KPC-2 | CTX-M-14 | SHV-28.v1 | OmpK35-21% | alive | yes |
| FQ39 | Other | SSI | 147 | rmp 1; KpVP-1 | rmpA2_6*-47% | ybt 9; ICEKp3 | iuc 1 |  |  | KL64 | NDM-1 | CTX-M-15 | SHV-11.v1 | OmpK35-25% | alive | yes |
| FQ148 | Pus | SSI | 231 | - | - | ybt 14; ICEKp5 | iuc unknown |  |  | KL51 | OXA-232 | - | SHV-1 | OmpK35-30% | alive | no |
| FQ162 | Urine | UTI | 231 | - | - | ybt 14; ICEKp5 | iuc unknown |  |  | KL51 | - | CTX-M-15 | - | OmpK35-30%; OmpK36TD | alive | yes |
| FQ62 | Blood | BSI | 231 | - | - | ybt 14; ICEKp5 | iuc unknown |  |  | KL51 | OXA-48 | CTX-M-14; CTX-M-15 | SHV-1 | OmpK35-30%; OmpK36-72% | died | no |
| FQ114 | RT | RTI | 231 | rmp 1; KpVP-1 | rmpA2_3*-47% | ybt 14; ICEKp5 | iuc unknown |  |  | KL51 | OXA-232 | CTX-M-15 | - | OmpK35-30%; OmpK36TD | died | no |
| FQ175 | RT | RTI | 231 | - | - | ybt 14; ICEKp5 | iuc unknown |  |  | KL51 | OXA-232 | CTX-M-15 | SHV-1 | OmpK35-30%; OmpK36TD | alive | yes |
| FQ185 | RT | RTI | 231 | - | - | ybt 14; ICEKp5 | iuc unknown |  |  | KL51 | - | CTX-M-15; SHV-106 | - | OmpK35-30%; OmpK36TD | alive | yes |
| FQ171 | Blood | BSI | 376 | rmp 1; KpVP-1 | - | - | iuc 1 |  |  | KL2 | OXA-48 | CTX-M-14 | - | - | died | yes |
| FQ168 | Urine | UTI | 383 | - | rmpA2_6*-47% | - | iuc 1 |  |  | KL30 | NDM-5; OXA-48 | CTX-M-14; CTX-M-15 | SHV-1 | OmpK35-10% | alive | no |
| FQ128 | Urine | UTI | 383 | rmp 1; KpVP-1 | rmpA2_6-60% | - | iuc 1 |  |  | KL30 | NDM-5; OXA-48 | CTX-M-14; CTX-M-15 | SHV-1 | OmpK35-10% | alive | no |
| FQ66 | Other | SSTI | 383 | rmp 1; KpVP-1 | rmpA2_6*-60% | - | iuc 1 |  |  | KL30 | OXA-48 | CTX-M-14 | SHV-1 | OmpK35-10% | alive | no |
| FQ61 | Pus | SSTI | 383 | rmp 1; KpVP-1 | rmpA2_6-60% | - | iuc 1 |  |  | KL30 | NDM-5; OXA-48 | CTX-M-14; CTX-M-15 | SHV-1 | OmpK35-10% | alive | no |
| FQ75 | Urine | UTI | 420 | rmp 1; KpVP-1 | rmpA2_3-47% | ybt 9; ICEKp3 | iuc 1 | *iro 1* |  | KL20 | OXA-48 | CTX-M-14 | SHV-75 | - | alive | no |
| FQ72 | RT | RTI | 420 | rmp 1; KpVP-1 | rmpA2_3-47% | ybt 9; ICEKp3 | iuc 1 | *iro 1* |  | KL20 | OXA-48 | CTX-M-14 | SHV-75 | - | alive | no |
| FQ48 | Urine | UTI | 2096 | - | rmpA2_8-60% | ybt 14; ICEKp5 | iuc 1 |  |  | KL64 | OXA-232; OXA-232 | CTX-M-15 | SHV-28.v1 | OmpK36GD | alive | yes |
| FQ156 | RT | RTI | ST25-1LV | - | - | ybt 14; ICEKp5 | - |  | clb 3 | KL2 | KPC-3; NDM-1 | CTX-M-15 | SHV-11.v1^ | OmpK36-50% | alive | no |
| FQ135 | Urine | UTI | ST35 | - | - | ybt 16; ICEKp12 | - |  |  | KL2 | OXA-181 | - | - | - | alive | no |
| FQ133 | Urine | UTI | ST39 | - | - | ybt 16; ICEKp12 | - |  |  | KL2 | NDM-1 | CTX-M-15 | SHV-11.v1^ | OmpK35-40%; OmpK36-55% | alive | no |

### Prevalence of virulence markers

Hypervirulence was defined as either (a) the presence of *rmpA* or *rmpA2*; and/or (b) the presence of aerobactin (*iuc*) and salmochelin (*iro*) (18). According to Kleborate results, 10 *K. pneumoniae* isolates (12.5%) had *rmpA1* and/or *rmpA2*, and three belonged to ST383. A combination of *iuc, iro* and *rmpA* were mostly concentrated together in ST147, ST383, ST420, and ST231 (Table 3). Sixteen (20%) *K. pneumoniae* isolates carried aerobactin (*iuc*), while only 2 (2.5%) isolates harboured salmochelin (*iro*), and they all belonged to ST420, and two (2.5%) isolates carried colibactin (*clb*) (one belonged to ST258). The yersiniabactin locus (*ybt* locus) encoding the acquired siderophore yersiniabactin was detected in 49 (61.3%) of *K. pneumoniae* and the most prevalent associated integrative conjugative elements (ICE*Kp*) were *ybt14* and ICE*Kp5*. All three *K. aerogenes* carried *iro*, and one of them (FQ126) also harboured *ybt* and *clb* (Table S1). The virulence loci *ybt* and *rmpA* were not detected in any of the study’s *K. oxytoca* and *K. quasipneumoniae* isolates. Capsule biosynthesis (K) loci were identified for all isolates, spanning 91 distinct K locus (KL) types (Table S1). Hypervirulent-associated *K. pneumoniae* serotypes KL1 and KL2 were rare; KL1 was not reported in this study, while KL2, usually associated with invasive liver abscess syndrome, was detected in five (5.8%) isolates of different STs, and one (FQ44) of them belonged to ST14. KL20 was detected in two ST420 isolates (Table 3).

### In-depth investigation Kp ST383

Out of four ST383 strains, two (FQ61, FQ128) carried both *rmpA* and *rmpA2*, as well as *bla*_NDM-5_ and *bla*_OXA-48_; in contrast FQ168 had both carbapenemases but did not have *rmpA*, while FQ66 had *rmpA/rmpA2* but did not have *bla*_NDM-5_. In order to understand the genetic basis and the plasmids associated with CR-hv *K. pneumoniae* in Qatar, we sequenced isolate FQ61 using both Illumina and MinION technologies. Hybrid assembly revealed that FQ61 harboured a chromosome and five plasmids, including pFQ61_ST383_NDM-5 (IncHI1B) (∼376kb), pFQ61_ST383_OXA-48 (IncL, ∼ 72kb), and Col (phAD28) (5-23kb) type plasmids. *bla*_*NDM*-5_ was located on the pFQ61_ST383_NDM-5 (IncHI1B type plasmidHow), which also carried eight other major AMR genes including *bla*_CTX-M-15_, *bla*_OXA-9_, *bla*_TEM-1_, *aac(6’)-Ib, aph(3’)-VI, aph(3’)-Ia, dfrA5, armA*. Based on BLAST search, a segment of this plasmid showed high similarity (>99%) with pKpvST383L (CP034201.2), a plasmid reported in another ST383 *K. pneumoniae* in the UK (19). Pairwise comparison revealed that the MDR region 1 (34,570 bp) was highly similar (99-100%) to the homologous region in pKpvST383L (Figure 2), while evidence of a large-scale inversion and rearrangement event was observed in the MDR region 2 adjacent to the repetitive elements (45,327bp) in comparison to pKpvST383L. The virulence region (42,300bp) harboring virulence genes *rmpA, rmpA2, iucABCD* and *iutA* also exhibited high similarity (> 99% identity and 100% query coverage) to pKvpvST383L (Figure 2), as well as pKpvST147B, pSI646A-ARMA-Vir-NDM, phvKpST395, phvKpST874, phvKpST147 in various *Klebsiella pneumoniae* isolates (20) (Figure 2). On the other hand, *bla*_OXA-48_ was located on the pFQ61_ST383_OXA-48 (IncL type plasmid), together with AMR genes such as *bla*_CTX-M-14b_ and *aph(3’)-VIb*. pFQ61_ST383_OXA-48 was highly similar (> 99.95%) to pKpvST383L (CP034202.1) in the UK, which was considered a hybrid virulence/resistance plasmid, carrying multiple insertional elements (IS), and multiple antimicrobial resistant and virulence genes (19).

**Figure 2.**
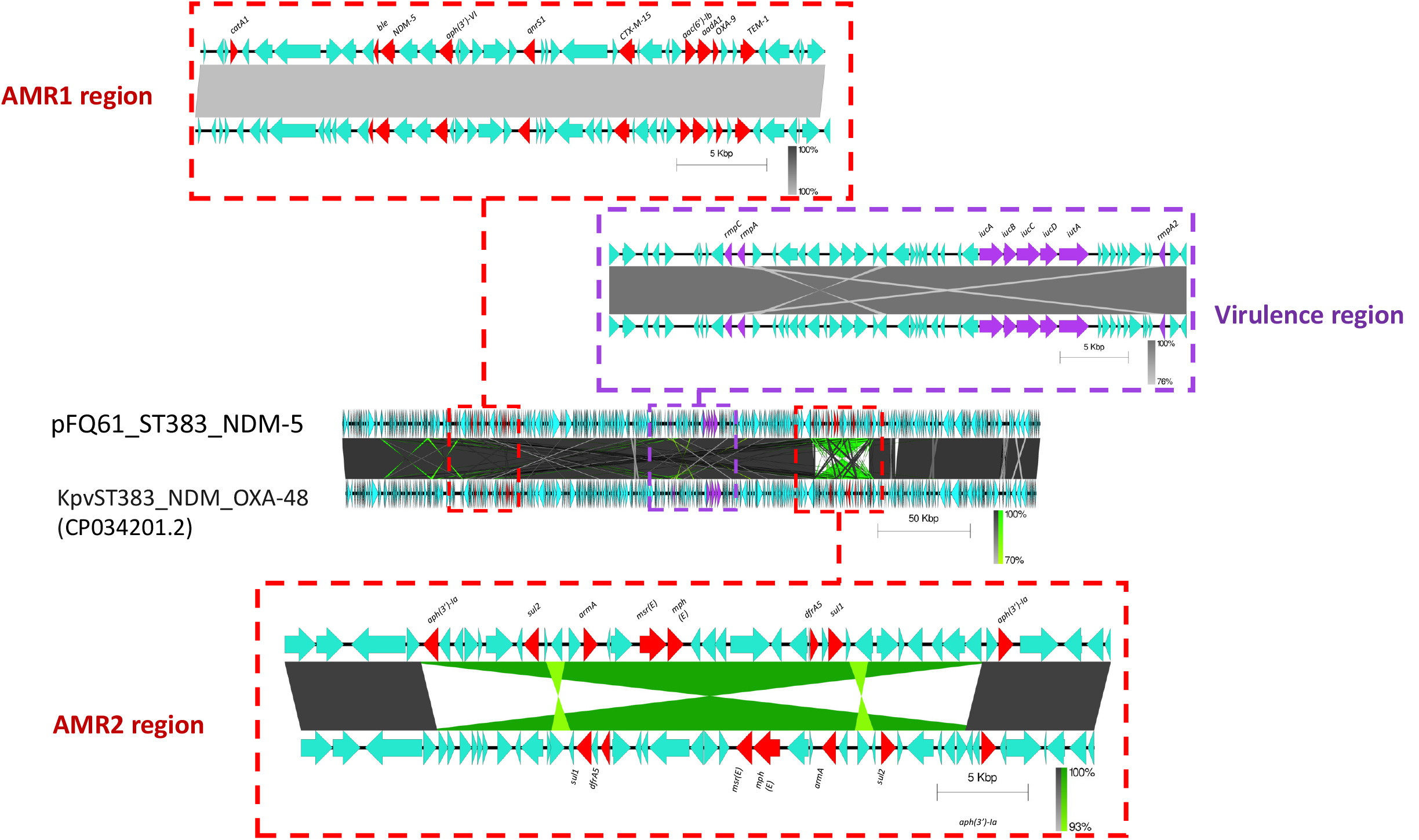
Genomic comparison of hybrid plasmid habouring *bla*_NDM-5_ and virulence phenotypes recovered from FQ61 and pKpvST383L[(19)].

Genomic comparison of our ST383 isolates with global ST383 isolates (Table S2) revealed the evolution of carbapenem-resistance and hypervirulence-associated genes in this particular lineage over the last decade. Figure 3 illustrates the phylogenetic relatedness of the local ST383 strains obtained together with publicly available ST383 assembled genomes and raw reads (N=32). The tree showed the isolates in Qatar (FQ61, FQ128, FQ168) clustered with those collected in Lebanon, the United Kingdom, and Italy, which also carried *bla*_NDM-5_/*bla*_NDM-1_, *bla*_OXA-48_, *bla*_CTX-M-15_, *bla*_CTX-M-14_, as well as several major virulence genes such as *iuc, rmpA1, rmpA2*. However, earlier reported ST383 isolates from Greece and France carried genes encoding carbapenemase such as KPC, OXA-48, and various VIM types (Figure 3), while isolates from China and Germany carried genes encoding OXA-48 only.

**Figure 3.**
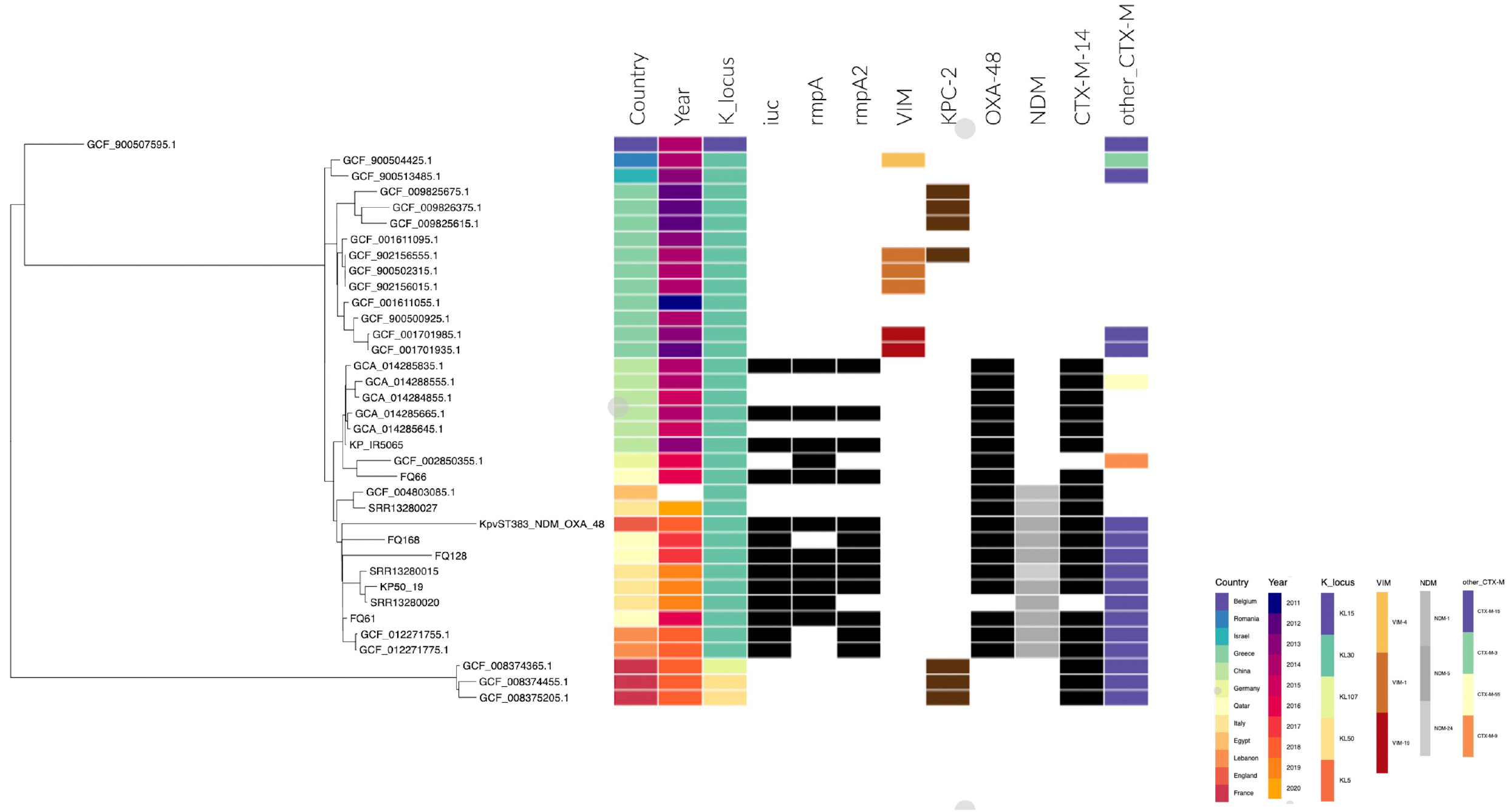
Phylogenetic tree showing the relationships among *K. pneumoniae* ST383 isolates from different countries.

We infected *G. mellonella* larvae with selected *K. pneumoniae* isolates. With an inoculum of 1.0×10^7^ CFU, the survival was 0% after 72 hours with classic hypervirulent K1 isolate (BL21, control), and 10% after 96 hours with two hypervirulent K2 isolates (FQ44, FQ156) (Figure 4). The ST383 isolates (FQ61, FQ128, FQ168) survival was 10-30% at 96 hours after infection. Also, the survival was 0% after 24 hours with FQ114 (ST231, *rmpA*+, *iuc*+) and 0% after 48 hours with FQ75 (ST420, *rmpA*+ *iuc*+ *iro*+ K20) and FQ179 (ST147, K64). All larvae injected with 10 mM MgSO_4_ solution only (negative control) survived (Figure 4).

**Figure 4.**
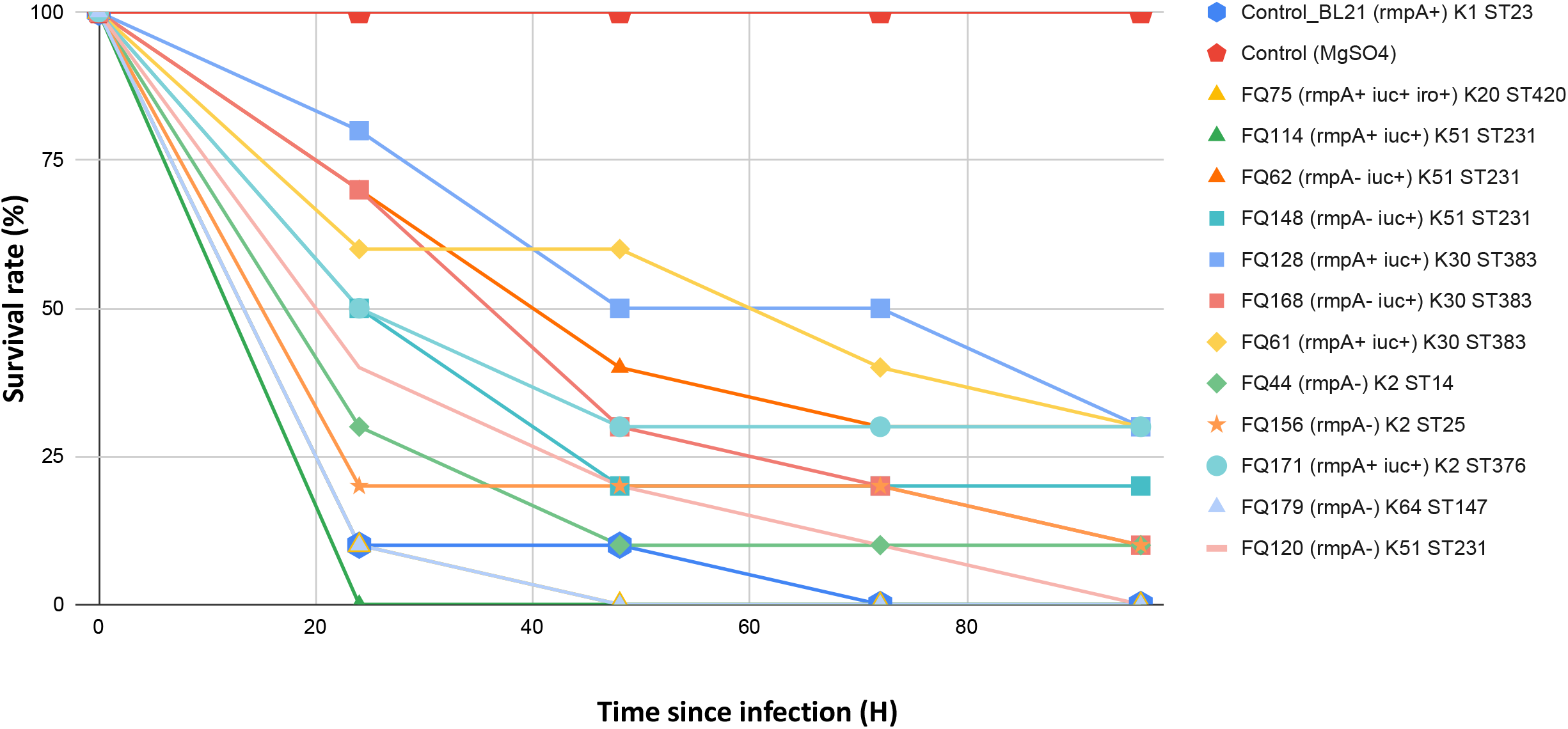
Virulence potential of selected *K. pneumoniae* isolates in a *Galleria mellonella* infection model.

### Phylogenetic analysis of *K. quasipnemoniae* var. *quasipnemoniae* ST196

WGS paired with epidemiological data clustered all *K. quasipnemoniae* var. *quasipnemoniae* ST196 isolates together, which prompted an investigation to identify possible disease outbreak (Figure 1). All nine ST196 isolates had identical MLST, capsular, and lipopolysaccharide types, and similar AMR genes (Table S1, Figures 1, 4), and the phylogenetic tree indicated they may represent a clone, as all isolates except FQ158 were highly similar (Figure 5). All *K. quasipnemoniae* var. *quasipnemoniae* ST196 were collected from various patients in different units, time, and departments of the same hospital system, which may represent hospital-acquired infections (Table S3).

**Figure 5.**
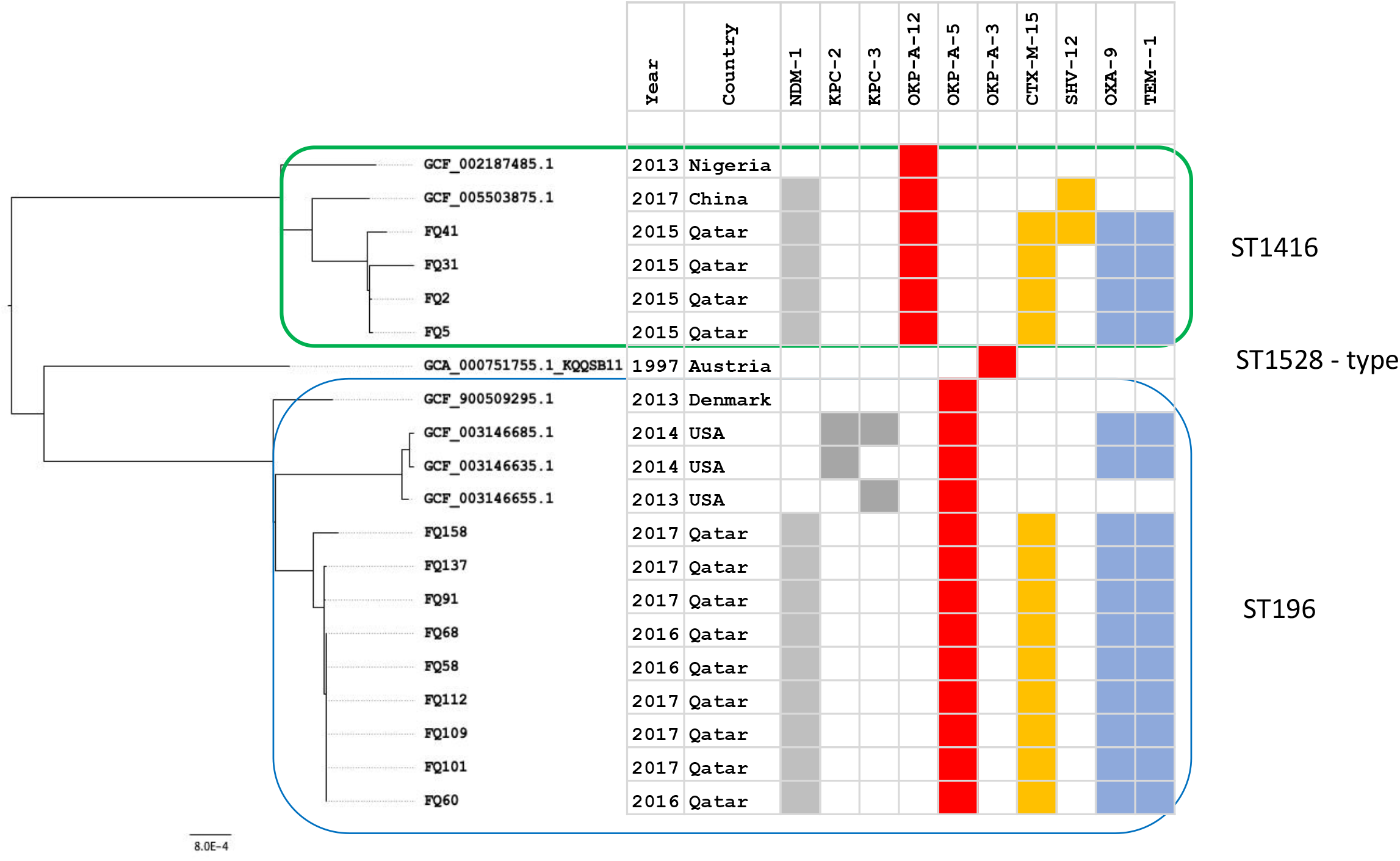
Phylogenetic relationships among *K. quasipneumoniae* ST196 and ST1416 isolates.

Most *K. quasipnemoniae* var. *quasipnemoniae* isolates in Qatar were NDM-1 producers, and the isolates belonging to ST196 were genetically different from global ST196 isolates (Figure 5). For instance, previously reported ST196 isolates were mostly KPC producers in Europe and the United States (Table S4). Virulence factors such as *iuc, clb, iro*, and *rmpA/rmpA2* genes were not detected in any of the *K. quasipnemoniae* var. *quasipnemoniae* isolates.

## Discussion

*Klebsiella pneumoniae* are common MDR pathogens in Qatar and are responsible for bloodstream and hospital-acquired infections (HAI). Our data revealed the presence of diverse STs and lineages in the region, and most were NDM-type and OXA-48-type producers co-haboring CTX-M ESBL genes. Among them, ST231, ST147 and ST11 were the most prevalent clones, which were different from those isolated from screening rectal swabs in local pediatric populations (7). ST147 (CG147) and ST11 (CG258) are international high-risk clones reported mainly from Asia and Europe and have been responsible for nosocomial transmission and various care-centre outbreaks (21, 22), while ST231 was considered an endemic clone disseminating OXA-232 in India (23). These STs could be obtained from various clinical specimens. *K. pneumoniae* CG258 (ST11 and ST258) is the predominant KPC-producing clone reported globally; however, only six isolates were reported in Qatar. International travel is an important mode of the spread of MDR bacteria, which may account for the diversity of carbapenem-resistant *Klebsiella* in Qatar and the Middle East region. In Qatar, substantial proportions of the population are migrant workers from the Indian subcontinent. Perez-Lopez (7) suggested that CRE in pediatric populations in Qatar were mainly introduced sporadically by asymptomatic carriers who visited or received health care in some nearby countries in which they are endemic. Moreover, consistent with other CRE studies on pediatric and adult patients, genes encoding NDMs and OXA-48 types were widely prevalent in Qatar, the Middle East region, and the Indian subcontinent (7, 22, 24). Few isolates were found to produce the KPC, VIM, or IMP, although these carbapenemases have been previously detected in Saudi Arabia and Kuwait (25).

Our results showed that classical hvKp, including CG23 and CG86, were rare among CR clinical isolates in Qatar. The proportion of hvKp among *K. pneumoniae* was 10% (*rmpA* and/or *rmpA2*), and can be up to 20% when solely aerobactin (*iuc*) bearing isolates were also included. The prevalence of *rmpA/rmpA2-* positive isolates was estimated to be 6% and 2.2% in Houston, USA and UK, respectively (26, 27). Also, the prevalence of hypervirulence was not that high and comparable to other countries, except China and Vietnam (> 20%) (10). *rmpA-* and/or *rmpA2*-mediated overproduction of capsular polysaccharide has been shown to contribute to hypervirulence (28), while *iuc* mediates increased siderophore production under iron-limiting conditions (16). In addition, KL1 and KL2 hypermucoviscous-specific capsular serotypes have been associated with invasive infections and accounted for approximately 70% of hvKp global isolates; however, they were not common in this collection. Low carriage of virulence genes among the CR isolates suggests that the circulation of hypervirulent strains such as ST23, ST65, and ST86 are uncommon in the clinical setting in this region. Convergence of carbapenem resistance and hypervirulence were found in limited isolates such as ST231, ST383 and ST410, of which ST231 and ST410 are globally emerging hypervirulent clonal groups associated with virulence plasmids (23, 29, 30).

Our molecular epidemiology data supported the persistence and emergence of a distinct local *K. quasipneumoniae var. quasipneumoniae* clone ST196. Indeed, *K. quasipneumoniae* ST196 and ST1416 carried *bla*_NDM-1_ and were different from isolates of the same STs reported in the US and Europe, which were usually KPC producers (https://doi.org/10.6084/m9.figshare.c.5238239.v1). Current data suggest the presence of an “endemic” NDM-1-producing clone in the Middle East region. The genomic similarity among these isolates collected from different patients/units was over 99%, and their MDR determinants and plasmid replicons were essentially identical. Outbreaks caused by *K. quasipneumoniae* have been rare, for instance ST334 has been reported as a potential emerging outbreak-associated MDR clone in Pakistan and Cambodia (31). Although the possibility of outbreak was ruled out, there may be unknown transmission mechanisms across facilities/sites or there may be ongoing circulation of this clone in the community, with our collection representing continued re-introduction into the hospital settings. In addition, hospitals can be the reservoir of these *K. quasipneumoniae* isolates and their plasmids, which may lead to persistence of clones and plasmids to cause additional infections and horizontal gene transfer.

Although the completeness of our assemblies and our abilities to identify plasmids are limited by the short read assemblies, our data indicated the dissemination of carbapenem resistance genes in *Klebsiella* was via acquisition of AMR plasmids. For instance, *bla*_OXA-48-like_ genes were often reported on the ColKP3 plasmids, particularly in ST147 and ST231 isolates. This is consistent with the report that the OXA48-like genes were carried mostly on Col plasmids (32). Also, *bla*_KPC-3_ in our collection was associated with the ‘traditional’ Tn*4401a* on the same contig, which can facilitate successful spread of KPC genes globally, particularly in North America.

Our study indicates the emergence of carbapenem-resistant and hypervirulent *K. pneumoniae* ST383 among patients in Qatar. Our patients infected with Kp ST383 had no travel histories, indicating they likely acquired the isolates locally. The wax moth larvae virulence study indicated that these ST383 isolates were able to cause mortality, even though our tested isolate (FQ61) was not as virulent as the classical hypervirulent KL1 isolate (control). Russo et al. (33) demonstrated that not all hvKp strains shared the same pathogenic potential in murine model infections. Previously, it has been common for *K. pneumoniae* ST383 isolates to be resistant to carbapenems, but not hypervirulent. Due to the acquisition of hybrid plasmid that contains a fraction of the hvKp virulence plasmid during the evolution of conventional ST383 isolates, these new circulating isolates have become both hypervirulent and multidrug-resistant, and should be regarded as a superbug that could pose a serious threat to the public health system. Early *K. pneumoniae* ST383 isolates carrying *bla*_VIM-4_, *bla*_KPC-2_, *bla*_CMY-2_ were reported in a Greek hospital during 2009-2010 (34). A later study demonstrated the presence of *bla*_VIM-19_ (35). More recent studies indicated the presence of *bla*_OXA-48_ in clinical isolates in Germany and China, which then went on to acquire another hybrid plasmid that carried both *bla*_NDM_ and virulence genes (19, 36), which represented a significant evolution event. Sabivora et al. argued the plasticity of the accessory genomes in ST383 isolate may benefit the acquisition of different plasmids (37); insertion elements could be responsible for mediating the plasmid recombination and rearrangement (14). Turton et. al. (26) studied 31 KL54 strains in the United Kingdom and identified 5 which harboured the *rmpA/rmpA2* and *iutA* genes. These ST383 isolates have also emerged in Italy recently (36, 38), and the Middle East region such as Qatar and Lebanon (Fig. 3). Recently, an ST383 isolate harbouring *bla*_KPC_ and genes encoding various virulence factors was reported in Saudi Arabia (39); however, assembled genome data are not available in the public domain. Tracking the evolution and distribution of ST383 is of major importance due to its ability to acquire carbapenemase genes of different types as well as genes associated with the hypervirulence phenotype. Increasing reports of the presence of hybrid, mosaic plasmid carrying both carbapenem resistance and virulence genes suggests CR-hvKp isolates are no longer confined to selected clones (14, 40), which will make containment of such isolates challenging.

In conclusion, our study has provided insights into the dynamics and epidemiology of *Klebsiella* in Qatar. Analysis of WGS data demonstrated the diversity of circulating STs and the presence of clonal lineage. Our comparative genomics data also confirmed the emergence of CR-hv Kp in Qatar, the Middle East region and worldwide. Acquisition of virulence genotype was reported in at least 10% CR Kp isolates, and one of the mechanisms was through the transfer of a carbapenem resistance and hypervirulence hybrid plasmid, as demonstrated in ST383 isolates, which are disseminating in different countries and continents. Molecular surveillance and monitoring of carbapenem-resistant *K. pneumoniae* isolates regarding AMR genes, and/or hybrid plasmids are important because isolates of convergent phenotypes would pose a health threat to the community and patients in health care facilities. Additional work using both short- and long-read data will facilitate identification of unique genetic and plasmid components for each species and clonal group. Further studies were required to understand the relationship between the hypervirulent phenotypes, carriage of the hybrid AMR-virulence plasmid and capsular types.

## Materials and Methods

### Cultures and antimicrobial susceptibility tests

As previously described (4), this study included carbapenem-resistant (CR) *Klebsiella* isolates from clinical specimens received at Hamad Medical Corporation’s Microbiology Department during the period between April 1, 2014 and November 30, 2017. The clinical data of the patients associated with the isolates were extracted from the electronic health records. National Healthcare Safety Network definitions were used to differentiate bacterial infection versus colonization. Matrix-Assisted Laser Desorption/Ionization-Time of Flight (MALDI-TOF) mass spectrometry (Bruker Corporation, Billerica, Massachusetts) was used for bacterial identification. The antimicrobial susceptibility testing for amikacin, cefotaxime, ceftazidime, ciprofloxacin, ertapenem, fosfomycin, gentamicin, meropenem, tigecycline and trimethoprim-sulfamethoxazole was performed on BD Phoenix™ (Becton, Dickinson and Company, Franklin Lakes, New Jersey, United States), using Clinical Laboratory Standards Institute breakpoints.^2^ *Enterobacterales* isolates were included if they were non-susceptible to ertapenem (minimum inhibitory concentration (MIC), >0.5 mg/L) or meropenem (MIC, >1.0 mg/L).^2^ Carbapenem resistance was confirmed using ertapenem disc diffusion method (zone of inhibition diameter of ≤21 mm).^2, 3^ The study was approved by the Institutional Review Board (MRC-16134/16).

### Whole genome sequencing (WGS) and data analysis

Genomic DNA was extracted using a DNeasy Blood and Tissue Kit (Qiagen, Hilden, Germany). DNA libraries were constructed with a modified Nextera XT DNA library preparation method (Illumina Inc., San Diego, California, USA), then sequenced on the Illumina NextSeq 550 platform with 2×150 cycles at Microbial Genome Sequencing Center (Pittsburgh, Pennsylvania, USA). The sequence data were assessed by Fastqc (http://www.bioinformatics.babraham.ac.uk/projects/fastqc/), trimmed by Trim Galore v0.6.0 (http://www.bioinformatics.babraham.ac.uk/projects/trim_galore/), assembled *de novo* using SPAdes v.3.9.0 (41) implemented in shovill (https://github.com/tseemann/shovill), and assessed using QUAST v5.0.2 (42). Contigs of smaller size (<500 bp), and with contaminants were excluded after analysis by Kraken v.2 (43). Sequence type (ST), plasmids and antimicrobial resistance genes were predicted from the assembled contigs using multi-locus sequence typing (MLST) (https://github.com/tseemann/mlst), the Plasmidfinder v2.1 (44) and ResFinder v3.2 databases implemented in ABRicate v0.9 (https://github.com/tseemann/abricate), based on >70% coverage and 90% sequence identity. Kleborate was used to detect the virulence genes, capsule synthesis (K) and lipopolysaccharide loci (O) (12).

Long reads were generated using an Oxford Nanopore MinION Sequencer (SQK-LSK109 and flow cell R9.4.1) at MiGS (Pittsburgh, USA). MinION reads were generated based on the Guppy software (v4.0.11) available from Oxford Nanopore Technologies. *De novo* Illumina-Nanopore assemblies were generated with Unicycler v0 · 4·7 (45). Genome assembly was annotated with Prokka v1·13·3 (46) and plasmid annotations were curated before depositing in GenBank. Phylogenetic trees were generated using Parsnp (47) and visualised together with associated metadata using Phandango v1.3 (48). Easyfig (49) was used to generate the diagram to compare the plasmid against the highly homologous plasmids in the NCBI database.

### Virulence study

The virulence of selected *K. pneumoniae* isolates, including three ST383, was tested in wax moth (*Galleria mellonella*) larvae. Briefly, overnight cultures of *K. pneumoniae* strains were prepared with 10 mM MgSO_4_ solution and further adjusted to concentrations of 1×10^5^ CFU/mL, 1×10^6^ CFU/mL, and 1×10^7^ CFU/mL. We infected the *G. mellonella* with the bacteria as described previously (50), and the survival rate of the *G. mellonella* was recorded every 24 h for 4 days.

## Data availability

Raw sequence reads are available on NCBI under BioProject accession number PRJNA656934. The genome sequence of FQ61 was submitted to GenBank under accession numbers CP091813-CP091818.

## Acknowledgements

The research was funded by the Medical Research Centre (MRC) at Hamad Medical Corporation (HMC) (MRC-16134/16 to F.B.) and supported by the National Institutes of Health (R01AI104895, R21AI135522, and R21AI151362 to Y.D.). We are grateful to clinical laboratory staff in the HMC, Dan Snyder at MiGS (Pittsburgh, USA), and staff in University of Pittsburgh for technical assistance. We also acknowledge Khalil Al Ismail, Head of Infection Control, Communicable Disease Center, Hamad Medical Corporation for performing outbreak investigation on *K. quasipneumoniae* ST196 isolates, Ebenezer Foster-Nyarko (Quadram Institute, UK) for assistance in plotting Figure 1, Kelly Wyres (Monash University, Australia) for advice, and Will Hsiao and Jun Duan (Simon Fraser University, Canada) for their technical support in bioinformatics analysis.

## Legends

Table 1. Comparison of key features of *Klebsiella* samples from different specimens in Qatar.

Table 2. Plasmid location of carbapenemases detected from the short read assemblies using Plasmidfinder.

Table 3. Notable hypervirulent *K. pneumoniae* isolates in this investigation.

## Supplementary materials

Table S1. A list of isolates with metadata, antimicrobial resistance genes and virulence phenotypes determined by Kleborate.

Table S2. The accession numbers of publicly available genomes/raw reads in KP ST383 used for comparative purposes

Table S3. Clinical information of ST196 *K. quasipneumoniae* isolates.

Table S4. The accession numbers of publicly available genomes ST1416 and ST196 *K. quasipneumoniae* isolates.

Figure S1. Diagram showing the genetic elements harbouring *bla*_KPC-2_ in *K. pneumoniae* isolates.

## References

1. Paczosa MK, Mecsas J. 2016. Klebsiella pneumoniae: Going on the Offense with a Strong Defense. Microbiol Mol Biol Rev 80:629–661.

2. Holt KE, Wertheim H, Zadoks RN, Baker S, Whitehouse CA, Dance D, Jenney A, Connor TR, Hsu LY, Severin J, Brisse S, Cao H, Wilksch J, Gorrie C, Schultz MB, Edwards DJ, Van Nguyen K, Nguyen TV, Dao TT, Mensink M, Minh VL, Nhu NTK, Schultsz C, Kuntaman K, Newton PN, Moore CE, Strugnell RA, Thomson NR. 2015. Genomic analysis of diversity, population structure, virulence, and antimicrobial resistance in Klebsiella pneumoniae, an urgent threat to public health. Proc Natl Acad Sci U S A 112:E3574–81.

3. Brisse S, Passet V, Grimont PAD. 2014. Description of Klebsiella quasipneumoniae sp. nov., isolated from human infections, with two subspecies, Klebsiella quasipneumoniae subsp. quasipneumoniae subsp. nov. and Klebsiella quasipneumoniae subsp. similipneumoniae subsp. nov., and demonstration that Klebsiella singaporensis is a junior heterotypic synonym of Klebsiella variicola. Int J Syst Evol Microbiol 64:3146–3152.

4. Abid FB, Tsui CKM, Doi Y, Deshmukh A, McElheny CL, Bachman WC, Fowler EL, Albishawi A, Mushtaq K, Ibrahim EB, Doiphode SH, Hamed MM, Almaslmani MA, Alkhal A, Butt AA, Omrani AS. 2021. Molecular characterization of clinical carbapenem-resistant Enterobacterales from Qatar. Eur J Clin Microbiol Infect Dis 40:1779–1785.

5. Baker S, Thomson N, Weill F-X, Holt KE. 2018. Genomic insights into the emergence and spread of antimicrobial-resistant bacterial pathogens. Science 360:733–738.

6. Al Mana H, Sundararaju S, Tsui CKM, Perez-Lopez A, Yassine H, Al Thani A, Al-Ansari K, Eltai NO. 2021. Whole-Genome Sequencing for Molecular Characterization of Carbapenem-Resistant Enterobacteriaceae Causing Lower Urinary Tract Infection among Pediatric Patients. Antibiotics (Basel) 10.

7. Pérez-López A, Sundararaju S, Tsui KM, Al-Mana H, Hasan MR, Suleiman M, Al Maslamani E, Imam O, Roscoe D, Tang P. 2021. Fecal Carriage and Molecular Characterization of Carbapenemase-Producing Enterobacterales in the Pediatric Population in Qatar. Microbiol Spectr 9:e0112221.

8. Perez-Lopez A, Sundararaju S, Al-Mana H, Tsui KM, Hasan MR, Suleiman M, Janahi M, Al Maslamani E, Tang P. 2020. Molecular Characterization of Extended-Spectrum β-Lactamase-Producing Escherichia coli and Klebsiella pneumoniae Among the Pediatric Population in Qatar. Front Microbiol 11:581711.

9. Jamshaid MB, Shahzad A, Zafar A, Kamal I. 2021. Invasive Klebsiella pneumoniae Syndrome in Qatar: A Case Report. Cureus 13:e15015.

10. Russo TA, Marr CM. 2019. Hypervirulent Klebsiella pneumoniae. Clin Microbiol Rev 32.

11. Russo TA, Olson R, MacDonald U, Beanan J, Davidson BA. 2015. Aerobactin, but not yersiniabactin, salmochelin, or enterobactin, enables the growth/survival of hypervirulent (hypermucoviscous) Klebsiella pneumoniae ex vivo and in vivo. Infect Immun 83:3325–3333.

12. Wyres KL, Nguyen TNT, Lam MMC, Judd LM, van Vinh Chau N, Dance DAB, Ip M, Karkey A, Ling CL, Miliya T, Newton PN, Lan NPH, Sengduangphachanh A, Turner P, Veeraraghavan B, Vinh PV, Vongsouvath M, Thomson NR, Baker S, Holt KE. 2020. Genomic surveillance for hypervirulence and multi-drug resistance in invasive Klebsiella pneumoniae from South and Southeast Asia. Genome Med 12:11.

13. Hennequin C, Robin F. 2016. Correlation between antimicrobial resistance and virulence in Klebsiella pneumoniae. Eur J Clin Microbiol Infect Dis 35:333–341.

14. Yang X, Dong N, Chan EW-C, Zhang R, Chen S. 2021. Carbapenem Resistance-Encoding and Virulence-Encoding Conjugative Plasmids in Klebsiella pneumoniae. Trends Microbiol 29:65–83.

15. Zhang R, Lin D, Chan EW-C, Gu D, Chen G-X, Chen S. 2016. Emergence of Carbapenem-Resistant Serotype K1 Hypervirulent Klebsiella pneumoniae Strains in China. Antimicrob Agents Chemother 60:709–711.

16. Wyres KL, Wick RR, Judd LM, Froumine R, Tokolyi A, Gorrie CL, Lam MMC, Duchêne S, Jenney A, Holt KE. 2019. Distinct evolutionary dynamics of horizontal gene transfer in drug resistant and virulent clones of Klebsiella pneumoniae. PLoS Genet 15:e1008114.

17. Wong JLC, Romano M, Kerry LE, Kwong H-S, Low W-W, Brett SJ, Clements A, Beis K, Frankel G. 2019. OmpK36-mediated Carbapenem resistance attenuates ST258 Klebsiella pneumoniae in vivo. Nat Commun 10:3957.

18. Lam MMC, Wyres KL, Duchêne S, Wick RR, Judd LM, Gan Y-H, Hoh C-H, Archuleta S, Molton JS, Kalimuddin S, Koh TH, Passet V, Brisse S, Holt KE. 2018. Population genomics of hypervirulent Klebsiella pneumoniae clonal-group 23 reveals early emergence and rapid global dissemination. Nat Commun 9:2703.

19. Turton J, Davies F, Turton J, Perry C, Payne Z, Pike R. 2019. Hybrid Resistance and Virulence Plasmids in “High-Risk” Clones of Klebsiella pneumoniae, Including Those Carrying blaNDM-5. Microorganisms 7.

20. Starkova P, Lazareva I, Avdeeva A, Sulian O, Likholetova D, Ageevets V, Lebedeva M, Gostev V, Sopova J, Sidorenko S. 2021. Emergence of Hybrid Resistance and Virulence Plasmids Harboring New Delhi Metallo-β-Lactamase in Klebsiella pneumoniae in Russia. Antibiotics (Basel) 10.

21. Navon-Venezia S, Kondratyeva K, Carattoli A. 2017. Klebsiella pneumoniae: a major worldwide source and shuttle for antibiotic resistance. FEMS Microbiol Rev 41:252–275.

22. Peirano G, Chen L, Kreiswirth BN, Pitout JDD. 2020. Emerging Antimicrobial-Resistant High-Risk Klebsiella pneumoniae Clones ST307 and ST147. Antimicrob Agents Chemother 64.

23. Shankar C, Mathur P, Venkatesan M, Pragasam AK, Anandan S, Khurana S, Veeraraghavan B. 2019. Rapidly disseminating blaOXA-232 carrying Klebsiella pneumoniae belonging to ST231 in India: multiple and varied mobile genetic elements. BMC Microbiol 19:137.

24. Zowawi HM, Sartor AL, Balkhy HH, Walsh TR, Al Johani SM, AlJindan RY, Alfaresi M, Ibrahim E, Al-Jardani A, Al-Abri S, Al Salman J, Dashti AA, Kutbi AH, Schlebusch S, Sidjabat HE, Paterson DL. 2014. Molecular characterization of carbapenemase-producing Escherichia coli and Klebsiella pneumoniae in the countries of the Gulf cooperation council: dominance of OXA-48 and NDM producers. Antimicrob Agents Chemother 58:3085–3090.

25. Alqahtani M, Tickler IA, Al Deesi Z, AlFouzan W, Al Jabri A, Al Jindan R, Al Johani S, Alkahtani SA, Al Kharusi A, Mokaddas E, Nabi A, Saeed N, Madian A, Whitmore J, Tenover FC. 2021. Molecular detection of carbapenem resistance genes in rectal swabs from patients in Gulf Cooperation Council hospitals. J Hosp Infect 112:96–103.

26. Turton JF, Payne Z, Coward A, Hopkins KL, Turton JA, Doumith M, Woodford N. 2018. Virulence genes in isolates of Klebsiella pneumoniae from the UK during 2016, including among carbapenemase gene-positive hypervirulent K1-ST23 and “non-hypervirulent” types ST147, ST15 and ST383. J Med Microbiol 67:118–128.

27. Chou A, Nuila RE, Franco LM, Stager CE, Atmar RL, Zechiedrich L. 2016. Prevalence of hypervirulent Klebsiella pneumoniae-associated genes rmpA and magA in two tertiary hospitals in Houston, TX, USA. J Med Microbiol 65:1047–1048.

28. Cheng HY, Chen YS, Wu CY, Chang HY, Lai YC, Peng HL. 2010. RmpA regulation of capsular polysaccharide biosynthesis in Klebsiella pneumoniae CG43. J Bacteriol 192:3144–3158.

29. Shankar C, Veeraraghavan B, Nabarro LEB, Ravi R, Ragupathi NKD, Rupali P. 2018. Whole genome analysis of hypervirulent Klebsiella pneumoniae isolates from community and hospital acquired bloodstream infection. BMC Microbiol 18:6.

30. Eger E, Heiden SE, Becker K, Rau A, Geisenhainer K, Idelevich EA, Schaufler K. 2021. Hypervirulent Klebsiella pneumoniae Sequence Type 420 with a Chromosomally Inserted Virulence Plasmid. Int J Mol Sci 22.

31. Wyres KL, Holt KE. 2016. Klebsiella pneumoniae Population Genomics and Antimicrobial-Resistant Clones. Trends Microbiol 24:944–956.

32. Li X, Ma W, Qin Q, Liu S, Ye L, Yang J, Li B. 2019. Nosocomial spread of OXA-232-producing Klebsiella pneumoniae ST15 in a teaching hospital, Shanghai, China. BMC Microbiol 19:235.

33. Russo TA, MacDonald U, Hassan S, Camanzo E, LeBreton F, Corey B, McGann P. 2021. An Assessment of Siderophore Production, Mucoviscosity, and Mouse Infection Models for Defining the Virulence Spectrum of Hypervirulent Klebsiella pneumoniae. mSphere 6.

34. Papagiannitsis CC, Giakkoupi P, Vatopoulos AC, Tryfinopoulou K, Miriagou V, Tzouvelekis LS. 2010. Emergence of Klebsiella pneumoniae of a novel sequence type (ST383) producing VIM-4, KPC-2 and CMY-4 β-lactamases. Int J Antimicrob Agents 36:573–574.

35. Papagiannitsis CC, Dolejska M, Izdebski R, Giakkoupi P, Skálová A, Chudějová K, Dobiasova H, Vatopoulos AC, Derde LPG, Bonten MJM, Gniadkowski M, Hrabák J. 2016. Characterisation of IncA/C2 plasmids carrying an In416-like integron with the blaVIM-19 gene from Klebsiella pneumoniae ST383 of Greek origin. Int J Antimicrob Agents 47:158–162.

36. Lorenzin G, Gona F, Battaglia S, Spitaleri A, Saluzzo F, Trovato A, di Marco F, Cichero P, Biancardi A, Nizzero P, Castiglione B, Scarpellini P, Moro M, Cirillo DM. 2022. Detection of NDM-1/5 and OXA-48 co-producing extensively drug-resistant hypervirulent Klebsiella pneumoniae in Northern Italy. J Glob Antimicrob Resist 28:146–150.

37. Sabirova JS, Xavier BB, Coppens J, Zarkotou O, Lammens C, Janssens L, Burggrave R, Wagner T, Goossens H, Malhotra-Kumar S. 2016. Whole-genome typing and characterization of blaVIM19-harbouring ST383 Klebsiella pneumoniae by PFGE, whole-genome mapping and WGS. J Antimicrob Chemother 71:1501–1509.

38. Spaziante M, Venditti C, Butera O, Messina F, Di Caro A, Tonziello G, Lanini S, Cataldo MA, Puro V. 2021. Importance of Surveillance of New Delhi Metallo-Beta-Lactamase Klebsiella pneumoniae: Molecular Characterization and Clonality of Strains Isolated in the Lazio Region, Italy. Infect Drug Resist 14:3659–3665.

39. Alghoribi MF, Binkhamis K, Alswaji AA, Alhijji A, Alsharidi A, Balkhy HH, Doumith M, Somily A. 2020. Genomic analysis of the first KPC-producing Klebsiella pneumoniae isolated from a patient in Riyadh: A new public health concern in Saudi Arabia. J Infect Public Health 13:647–650.

40. Huang Y-H, Chou S-H, Liang S-W, Ni C-E, Lin Y-T, Huang Y-W, Yang T-C. 2018. Emergence of an XDR and carbapenemase-producing hypervirulent Klebsiella pneumoniae strain in Taiwan. J Antimicrob Chemother 73:2039–2046.

41. Bankevich A, Nurk S, Antipov D, Gurevich AA, Dvorkin M, Kulikov AS, Lesin VM, Nikolenko SI, Pham S, Prjibelski AD, Pyshkin AV, Sirotkin AV, Vyahhi N, Tesler G, Alekseyev MA, Pevzner PA. 2012. SPAdes: a new genome assembly algorithm and its applications to single-cell sequencing. J Comput Biol 19:455–477.

42. Gurevich A, Saveliev V, Vyahhi N, Tesler G. 2013. QUAST: quality assessment tool for genome assemblies. Bioinformatics 29:1072–1075.

43. Wood DE, Salzberg SL. 2014. Kraken: ultrafast metagenomic sequence classification using exact alignments. Genome Biol 15:R46.

44. Carattoli A, Zankari E, García-Fernández A, Voldby Larsen M, Lund O, Villa L, Møller Aarestrup F, Hasman H. 2014. In silico detection and typing of plasmids using PlasmidFinder and plasmid multilocus sequence typing. Antimicrob Agents Chemother 58:3895–3903.

45. Wick RR, Judd LM, Gorrie CL, Holt KE. 2017. Unicycler: Resolving bacterial genome assemblies from short and long sequencing reads. PLoS Comput Biol 13:e1005595.

46. Seemann T. 2014. Prokka: rapid prokaryotic genome annotation. Bioinformatics 30:2068–2069.

47. Treangen TJ, Ondov BD, Koren S, Phillippy AM. 2014. The Harvest suite for rapid core-genome alignment and visualization of thousands of intraspecific microbial genomes. Genome Biol 15:524.

48. Hadfield J, Croucher NJ, Goater RJ, Abudahab K, Aanensen DM, Harris SR. 2018. Phandango: an interactive viewer for bacterial population genomics. Bioinformatics 34:292–293.

49. Sullivan MJ, Petty NK, Beatson SA. 2011. Easyfig: a genome comparison visualizer. Bioinformatics 27:1009–1010.

50. O’Hara JA, Ambe LA, Casella LG, Townsend BM, Pelletier MR, Ernst RK, Shanks RMQ, Doi Y. 2013. Activities of vancomycin-containing regimens against colistin-resistant Acinetobacter baumannii clinical strains. Antimicrob Agents Chemother 57:2103–2108.

